# VeloSim: Simulating single cell gene-expression and RNA velocity

**DOI:** 10.1101/2021.01.11.426277

**Authors:** Ziqi Zhang, Xiuwei Zhang

## Abstract

The availability of high throughput single-cell RNA-Sequencing data allows researchers to study the molecular mechanisms that drive the temporal dynamics of cells during differentiation or development. Recent computational methods that build upon single-cell sequencing technology, such as trajectory inference or RNA-velocity estimation, provide a way for researchers to analyze the state of each cell during a continuous dynamic process. However, with the surge of such computational methods, there is still a lack of simulators that can model the cell temporal dynamics, and provide ground truth data to benchmark the computational methods.

Hereby we present `VeloSim`, a simulation software that can simulate the gene-expression kinetics in cells along continuous trajectories. `VeloSim` is able to take any trajectory structure composed of basic elements including “linear” and “cycle” as input, and outputs unspliced mRNA count matrix, spliced mRNA count matrix, cell pseudo-time and true RNA velocity of the cells. We demonstrate how `VeloSim` can be used to benchmark trajectory inference and RNA-velocity estimation methods with different amounts of biological and technical variation within the datasets. `VeloSim` is implemented into an R package available at https://github.com/PeterZZQ/VeloSim.

## 1 Introduction

Applying Single-cell RNA-Sequencing (scRNA-Seq) on cell populations during cell differentiation or development helps researchers to capture each individual cell from different developmental stages within the population at the same time. Computational methods such as trajectory inference^1–3^ and RNA-velocity^4, 5^ are used to recover the states of cells and infer the trajectory of the cell dynamic processes in the population, using scRNA-seq data. Trajectory inference reconstruct the trajectory structure in the dataset, assign cells onto different trajectory paths and order cells according to their developmental stages. RNA-velocity, on the other hand, provides a short term prediction of gene expression profile for each cell using unspliced and spliced mRNA count. However, it is a challenging problem to learn the correct trajectories or to predict future states from scRNA-seq data due to the high amount of noise in the data and the complex nature of cell dynamics.

Simulated datasets have been used to benchmark the trajectory inference methods and learn the strength and weakness of each method^6, 7^. However, there are less existing benchmarking work for methods which involve RNA velocity, as most of the simulators, including Splatter^8^, SymSim^7^, powSimR^9^. Dyngen^10^ and SERGIO^11^ are existing methods which able to simulate both spliced and unspliced mRNA counts of cells in a continuous dynamic process, with an underlying gene regulatory network.

We present `VeloSim`, an R package that can simulate both unspliced and spliced mRNA counts of single cells along continuous trajectories, as well as the true RNA velocity. It also outputs the assignment of cells to each trajectory lineage, and the pseudo-time of each cell. To simulate the unspliced and spliced mRNA counts, `VeloSim` uses the two-state kinetic model, where a gene is considered to be in an *on* or *off* state, and when in an *on* state, the unspliced mRNAs are being produced, and whenever there are unspliced mRNAs, spliced mRNAs are produced using the unspliced mRNAs. `VeloSim` allows users to easily provide any trajectory structure that is made of basic elements of “cycle” and “linear”. A tree structure can be generated simply by combining multiple linear structures. `VeloSim` can be used to benchmark different computational methods designed for continuous developmental dataset including trajectory inference and RNA-velocity estimation. The schematics of `VeloSim` is shown in Fig. 1.

**Figure 1.**
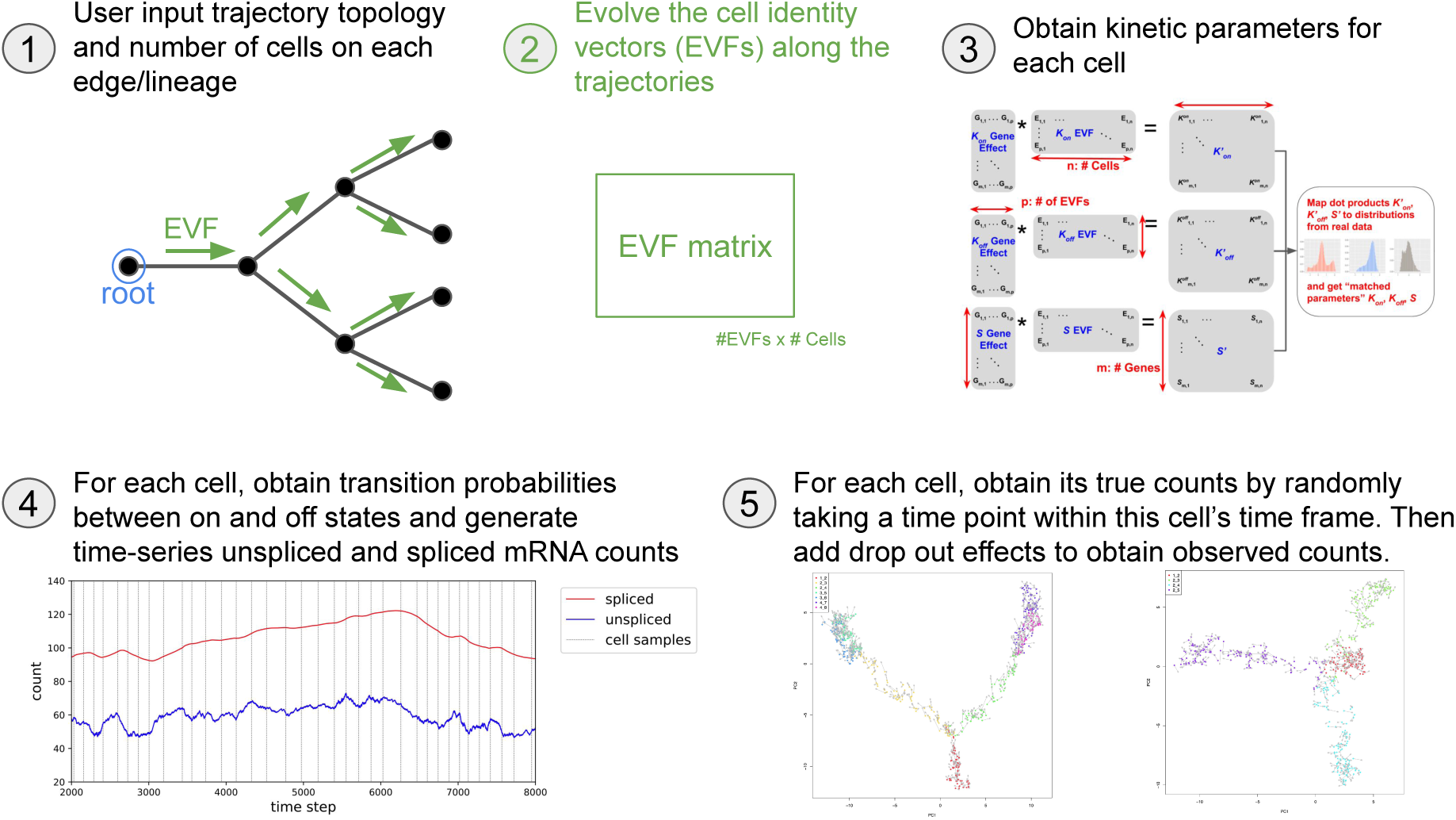
The schematics of `VeloSim`. `VeloSim` first samples cell extrinsic variability factors(EVFs) from the trajectory topology, and generates the kinetic parameters using gene effect matrix and EVF matrix. With simulated kinetic parameters, `VeloSim` generates the time-series unspliced and spliced mRNA counts using the kinetic model. Finally, true counts of each cells are sampled from the time-series counts data, and the observed counts are generated from the true counts by adding in dropout effect.

## 2 Results

### 2.1 VeloSim models time-series gene expression using the kinetic model

Following SymSim^7^, `VeloSim` models the mRNA transcription process with two-state kinetic model^12, 13^, which incorporates on state and *off* state of a gene in a cell. The mRNAs are transcribed if the corresponding gene is at the *on* state, and are inhibited if the gene is at the *off* state. The transition of each gene between those two states are modeled by a two state markov process, with the transition probability calculated from *k*_on_, the rate at which the a gene becomes active, and *k*_off_, the rate at which the gene is inhibited. We generalized the kinetic model used in^7^ and^13^ to further incorporate the splicing process of mRNA, in order to record the unspliced mRNA counts. The mRNA molecules before the splicing process is called nascent mRNA or unspliced mRNA, and after the process is called mature mRNA or spliced mRNA. The whole process can be modeled using a group of first order differential equation (ODE) (Methods)^4, 14^ with additional parameters *s, β* and *γ* that correspond to transcription rate, splicing rate and degradation rate of the mRNA molecules. Similar modeling was used in the original RNA velocity paper by La Manno *et al*^4^.

To introduce how `VeloSim` generates time-series unspliced counts and spliced counts for a given gene along a trajectory lineage, we assume that we know the kinetic parameters, *k*_on_, *k*_off_, *s, β* and *γ*, of this gene’s expression level in these cells. Then for a gene in each cell, we can generate how its unspliced and spliced counts change for a period of time (Methods), then we move on to the expression of this gene in the next cell. Fig. 2 shows the change of unspliced and spliced counts of a gene in a sequence of cells, with different resolutions. To mimic the snapshot property of scRNA-seq data, for each cell, we randomly select a time point within its time range, and take the value of unspliced and spliced counts for all genes at this time point, as the true counts for all genes in this cell.

**Figure 2.**
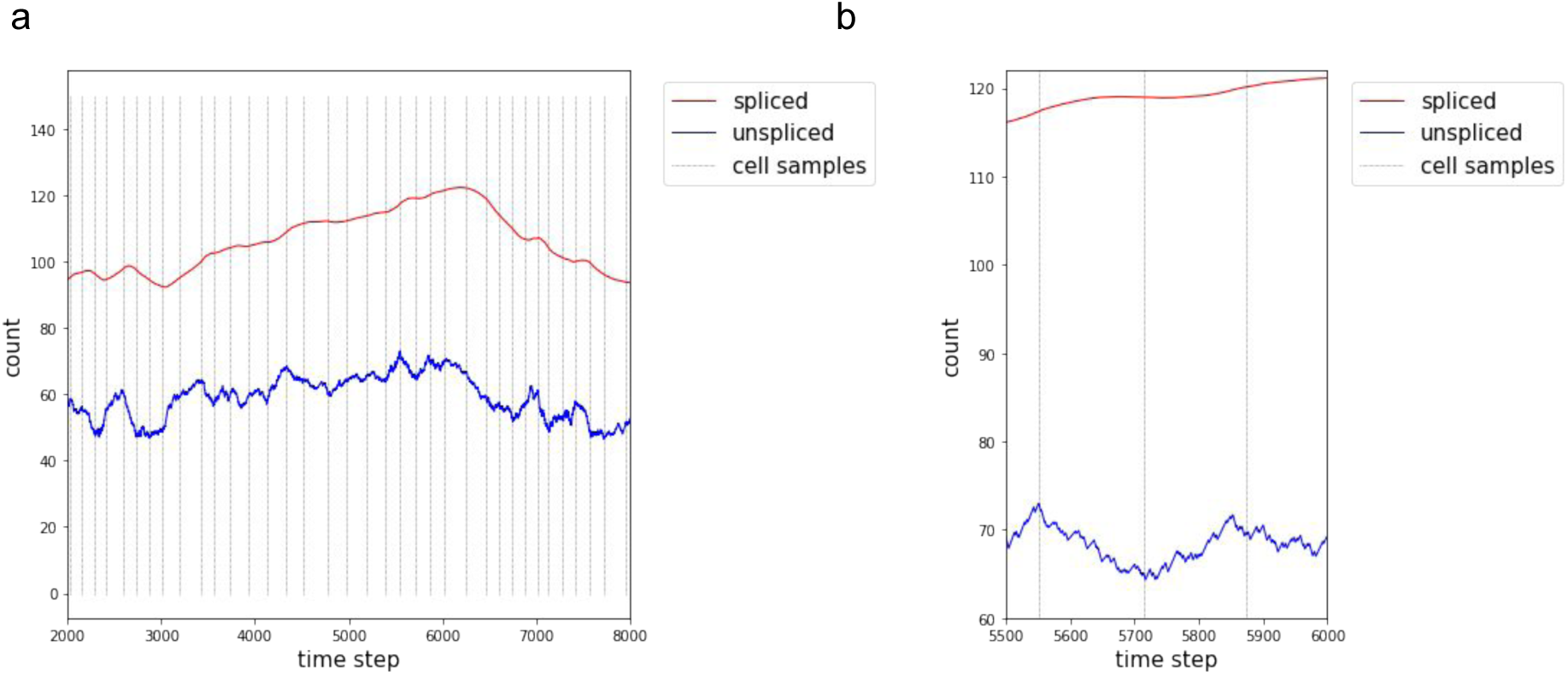
The dynamics of unspliced and spliced mRNA count during cell differentiation, simulated with `VeloSim`. Every two consecutive cells are separated by a vertical gray line. (b) shows a zoomed in version of part of (a).

### 2.2 VeloSim models the gradual change of cell identities using EVFs

The kinetic parameters used to generate the dynamic process described above are generated based the cell identities and gene identities. We assume that *k*_on_, *k*_off_ and s are generated differently for each gene in each cell, and the splicing rate *β* and degradation rate *γ*, are gene-specific parameters and are shared between cells. The kinetic parameters *k*_on_, *k*_off_ and s are estimated using the dot product of cell identity vectors and gene identity vectors. *β* and *γ* are sampled independent and identically from normal distribution with user-defined mean and variance.

Cell identity vectors, or we termed extrinsic variability factors(EVF)s, are low dimensional vectors unique for each cell. It serves as a latent representation of each cell within the cell population, which can models multiple biological factors, such as morphology, maturity, functionality and micro-environment. We dichotomize the factors into two subsets, one include the the factors that is driving the differentiation, termed differential EVF, another include the factors that is similar for all cells within the cell population, termed non-differential EVF. Totally three vectors are generated for each cell, with one vector for each kinetic parameter: *k_on_, k_off_* and *s*. In order to simulate a continuous cell population where cells from different developmental stages are incorporated, we generate differential EVF using the Brownian motion procedure along the differentiating backbone^15^, such as tree or cycle. The non-differential EVFs, on the other hand, are sampled directly from an i.i.d normal distribution with mean and variance provided by the user.

The gene identity vectors, also called gene effects, are vectors of the same length as the EVF vectors, and for a given gene and a given cell, the gene effect vector represent how much this gene is affected by different EVFs in the EVF vector of the cell. In `VeloSim`, we first set a majority of values in the gene effect vectors to be 0, then generate the non-zero values from a Gaussian distribution. In order to produce a more realistic kinetic estimation, we further match the parameter distribution of the inferred kinetic to the kinetic from real world dataset with quantile normalization.

With all the kinetic parameters being estimated, we generate the unspliced and spliced mRNA count using differential equations that model the gene expression dynamics(Fig. 2). We generate one linear trajectory with one simulation on the equations given the starting states and kinetic parameters. The complex tree lineage topology is generated by running the linear trajectory generation process for multiple times with different kinetic parameters for the lineage and starting states extracted from the ending states of previous linear trajectory segment. For cycle topology, the generation process is exactly the same as the generation process of linear trajectory, but we use a different Brownian motion process such that to generate the EVFs such that cells at the end of the cycle have similar identity as the cells at the beginning of the cycle (Methods). We dichotomies the kinetic parameter generation process into forward and backward parts, where cell develops forward to the turning point and backward to the original state. The generated sample scRNA-Seq data of binary-tree topology and cycle-topology is shown in Fig. 3.

**Figure 3.**
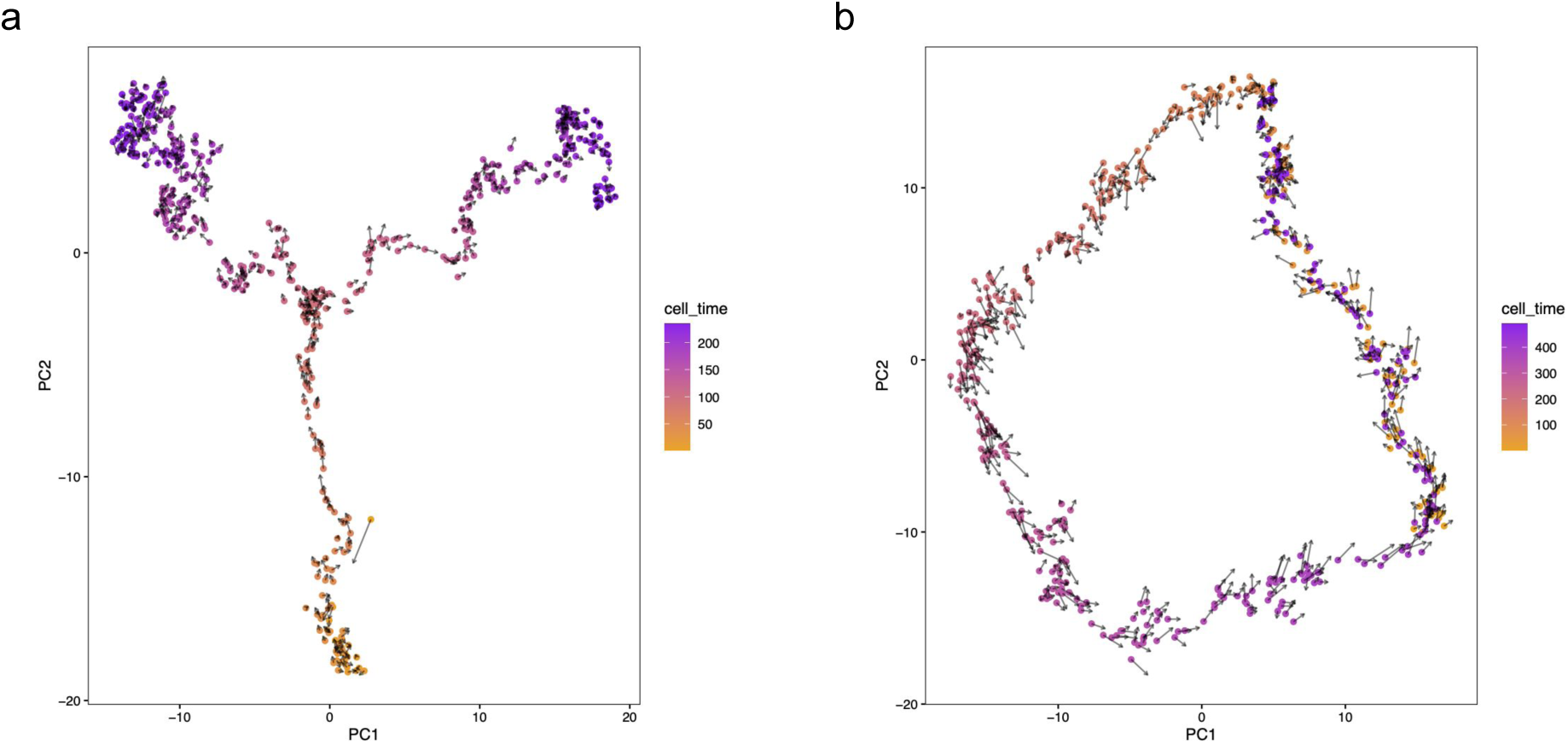
(a). PCA visualization of scRNA-Seq dataset generated by `VeloSim` with binary tree topology. Cells are colored with cell developmental pseudo-time and the arrow denotes the velocity direction. (b). PCA visualization of scRNA-Seq dataset generate by `VeloSim` with cell cycle topology.

In addition to simple cycle and tree topology, `VeloSim` is able to generate the complex cycle-tree structured dataset, where cell-cycle process coexists with the tree-like cell differentiation process^16^. Such trajectory structure exists in real life dataset^5^, but there are limited simulators that can simulate such trajectory structure. `VeloSim` generate the structure by first simulating the kinetic parameters a multi-cycle cell cycling process, then a tree like branching process. The parameter distribution of the inferred kinetic are matched to the kinetic from real world dataset jointly, and cells in the topology are generated using the inferred kinetic parameters.

After the count matrix is generated, `VeloSim` simulate the dropout effect of real life sequencing process by sampling the count matrix with small molecule capture rate (Methods).

### 2.3 Using VeloSim to benchmark trajectory inference methods

`VeloSim` can be used to benchmark different trajectory inference methods that require either root cell or RNA-velocity information. We benchmark `slingshot`^2^ and `latent` time^5^ methods, one requires root cell information, another uses RNA velocity information. We provide ground truth root cell cluster for `slingshot` and RNA velocity information for `latent time`. We test both methods on multiple dataset with tree-like topology, and measure the performance of different methods using kendall rank correlation coefficient (Methods), the result (Fig. 4) shows that if given correct root cell, `slingshot` generally performs better compared to `latent time` in tree-like trajectory structure.

**Figure 4.**
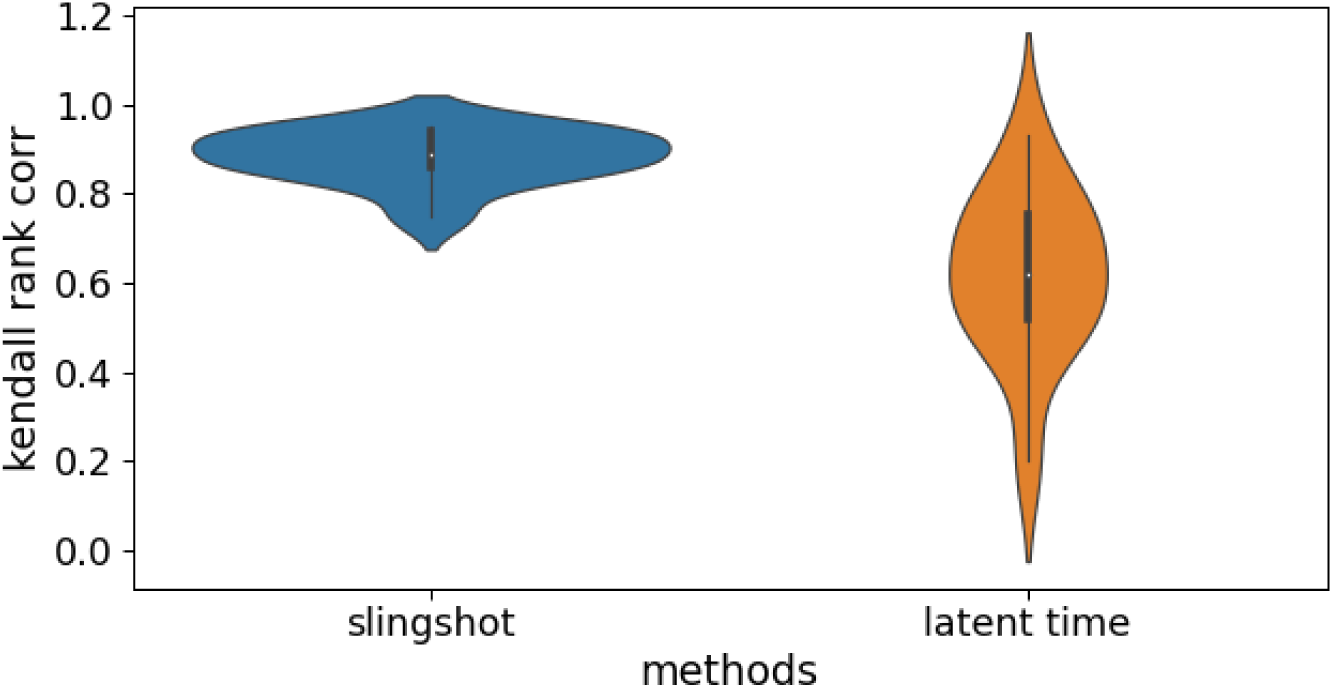
Pseudo-time inference accuracy calculated with `slingshot` and latent time, the kendall rank correlation coefficient is measured between inferred pseudo-time and ground truth pseudo-time. Slingshot perform generally better than latent time, but requires the root cell is provided in advance.

### 2.4 Using veloSim to benchmark RNA-velocity estimation methods

Providing unspliced, spliced and true RNA-velocity count matrix at the same time, `VeloSim` is able to benchmark different RNA-velocity inference methods easily. We use `VeloSim` to benchmark different modes of velocyto^4^ and `scVelo`^5^. We use a bifurcating trajectory structure, and generated 27 datasets with different parameters including random seed, number of cells, number of genes. We test the performance of different RNA velocity inference methods with different molecular capture rate, which correspond to the level of dropout effect induced by real-life sequencing process, and the result(Fig. 5) shows that all current RNA velocity inference methods are sensitive to dropout effect within the dataset, and the stochastic mode of `scVelo` outperform all the other methods under all scenarios with different capture rates. The RNA velocity inference accuracy of each cell is measured using cosine similarity (Methods).

**Figure 5.**
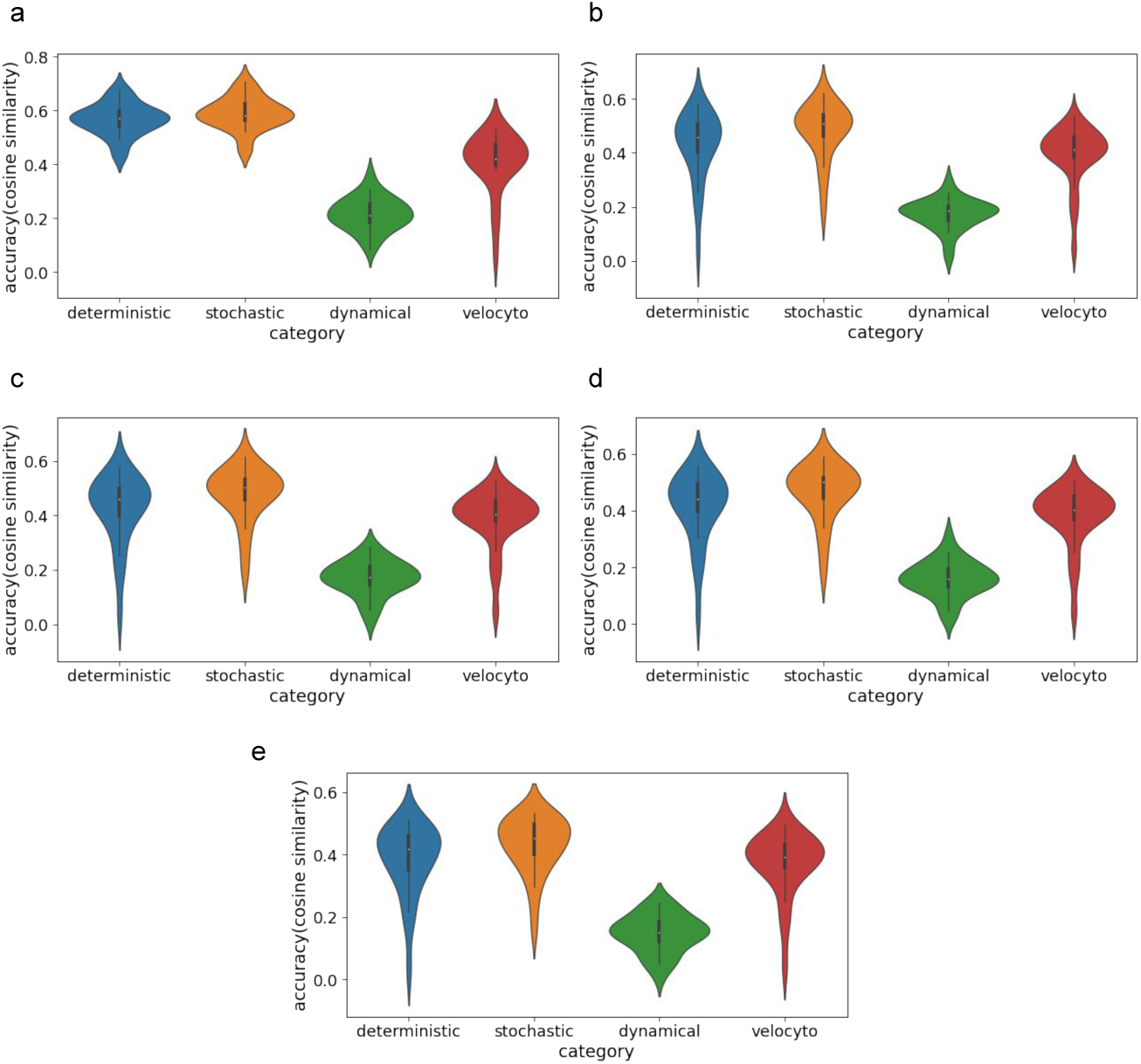
(a). The cosine similarity score of RNA velocity inferred with different methods, using count matrix with capture rate *c* = 1. (b). The score of RNA velocity inferred with different methods, using count matrix with capture rate *c* = 0.8. (c). The score of RNA velocity inferred with different methods, using count matrix with capture rate *c* = 0.6. (d). The score of RNA velocity inferred with different methods, using count matrix with capture rate *c* = 0.4. (e). The score of RNA velocity inferred with different methods, using count matrix with capture rate *c* = 0.2.

## 3 Discussion

We proposed `VeloSim`, an R package that generates unspliced and spliced mRNA counts for single cells along any continuous trajectories. It is an easy-to-use package which can generate ground truth trajectory topology and cell pseudo-time, and can be used to benchmark trajectory inference methods and RNA velocity inference methods.

## 4 Methods

### 4.1 Generating time-series unspliced and spliced counts using kinetic model

In^4^, the authors modeled the generation of unspliced and spliced counts as follows:

Denoting the transcription rate as *α*, the amount of unspliced mRNA as *u*, the splicing rate (the rate of unspliced mRNA turning into spliced mRNA) as *s*, and the degradation rate as *γ*, then we have

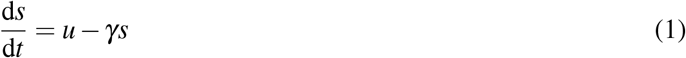

In `VeloSim`, we know the kinetic parameters *k*_on_, *k*_off_ and *s*. 1/*k*_on_ represents the average waiting time of the gene in the *off* state before it is switched on; 1/*k*_off_ represents the average waiting time of the gene in the *on* state before it is switched off. We aim to calculate the transition probability between *on* and *off* states using the *k*_on_ and *k*_off_. First, we define a *cycle* of a gene as *T_c_* = 1/*k*_on_ + 1/*k*_off_. Then we divide this time window into *n*_part_ parts, where *n*_part_ = max{*T_c_*/ min(1/*k*_on_, 1/*k*_off_), *T_c_*} * 2, then each part has length Δ *t* = *T_c_*/*n*_part_, and this is also what we call *stepsize*. With this stepsize length, the transition probability matrix *P*_tran_ between the on and off states can be calculated as:

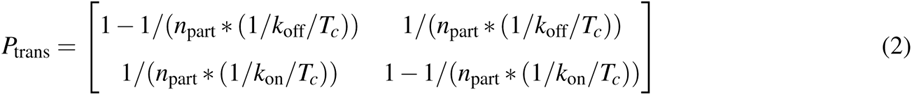

where *P*_tran_ [1, 1] represents the probability of going to an on state from an on state after time of stepsize, *P*_tran_ [1, 1] represents the probability of going to an off state from an *on* state after time of stepsize, etc.

We start with assigning the gene a random state (*on* or *off*), then simulate its gene-expression in terms both unspliced and spliced counts from cell to cell. For each cell, we run the time of a full cycle *T_c_*, depending on the kinetic parameters of the gene in this cell. At every step (with stepsize Δ*t*), we apply the transition probability matrix *P*_tran_ to determine the next state. If the gene is now in an *on* state at time *t*, the unspliced counts at time *t* is calculated as:

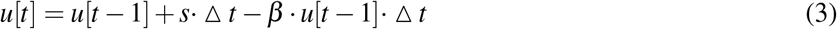

If at time *t*, the gene is in an off state, we have:

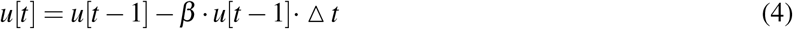

Then we update the amount of spliced counts:

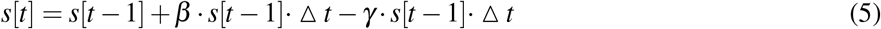

In this fashion, we can simulate the unspliced and spliced counts over a time course of *T_c_* for each cell. Then, a random time point *t_r_* is selected during the time course of *T_c_* to extract the corresponding unspliced and spliced counts at time *t_r_*, as the data for this gene in this cell. This mimics the snapshot property of single cell RNA-seq data.

### 4.2 Cell EVF and gene effect vector generation

The cell EVFs are generated along the given trajectory structure with a Brownian motion process, as used in SymSim^7^. This process mimics that the cells’ identities change gradually along a continuous trajectory. This framework also allows users to input any trajectory, consisting of basic components like edges in a tree and cycles.

Generating a cycle structure is a bit more tricky. In a cycle structure, we need to ensure that the last cell “connect” to the first cell. To realize this, we use the following strategy. Suppose we would like to generate *n_c_* cells along a cycle structure. First we set *e*_1_ = 1, then for cells 2 to ⌈*n_c_*/2⌉, we generate the EVF values using the standard process, which is

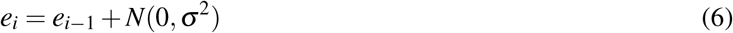

where *e_i_* represents the EVF value of cell *i*, and *N*(0, σ^2^) means a value sampled from a Gaussian distribution with mean 0 and standard deviation σ. σ is a parameter which can adjust how close the cells are to the backbone trajectory structure. Smaller σ makes the cells deviates less from the backbone structure.

While we perform Eq. 6 for *i* = 2,…, ⌈*n_c_*/2⌉, we record the value sampled from *N*(0, σ^2^) and set the corresponding values as *d*_2_,…, *d*_⌈*n_c_*/2⌉_. Now to generate *e_j_* for *k* = ⌈*n_c_*/2⌉ +,…,*n_c_*, for every *j*, we randomly select a value from *d*_2_,…,*d*_⌈*n_c_*/2⌉_ (withoutreplacement), say we select *d_k_*, then we do:

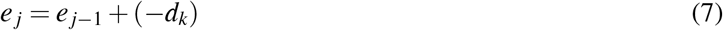

In this fashion, we can ensure that *e_k_* is very close or equal to *e*_1_, with small difference (sampled from *N*(0, σ^2^)) between any two consecutive EVF values.

Gene effect vectors represent the identity of genes. In `VeloSim` we do not incorporate gene regulatory networks or coexpression gene modules, so the gene effect vectors of genes are generated in the same way: in each gene effect vector, we first set certain positions to be 0s, and for the non-zero entries, we sample

We also models the highly expressed genes that does not follows the general kinetic value distribution. We define a user-specified parameter *prop_hge_*, which correspond to the proportion of the highly expressed genes, and adjust the transcription rate s for those genes to increase their final count.

### 4.3 Adding technical noise to unspliced and spliced counts

Real life count matrix obtained from single-cell RNA Sequencing technology usually exhibit high sparsity due to strong molecular dropout effect. We simulate the sequencing process by sampling the true mRNA count with molecular capture probability *c*(*c* = 0.2). With each mRNA molecule, there is a probability *c* = 0.2 that this molecule is captured in the sequenced count.

### 4.4 Comparing different RNA velocity inference methods

We test the performance of different RNA velocity inference methods with multiple datasets generated with different kinetic parameters. We generate totally 27 single-cell datasets with number of cells set to 500, 750, 1000, number of genes set to 100, 200, 500, and random seeds set to 2, 3 and 4. With different each dataset, we test how the performance of RNA velocity inference varies under different capture rate (*c* = 0.2, 0.4, 0.6, 0.8 and 1). Totally four different methods are tested, including `velocyto, scVelo-deterministic` mode, `scVelo-stochastic` mode and `scVelo-dynamical` mode. The inference accuracy is measured using cosine similarity, which measures the angle between two inferred velocity directions. For cell *i*, denote the ground truth RNA velocity as vector **v**_*t*_(*j*) and the inference RNA velocity as vector **v**_*i*_(*j*), and the cosine similarity can be calculated as

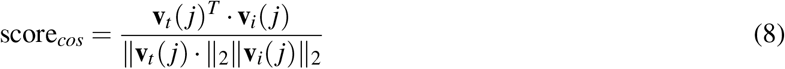

The score lies within the range between –1 and 1, with higher score correspond to the case where the inferred velocity direction is highly similar to the ground truth velocity direction.

### 4.5 Comparing different trajectory inference methods

We measure the pseudo-time inference accuracy using kendall rank correlation coefficient between inferred pseudo-time and ground truth pseudo-time. Given 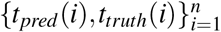, the kendall rank correlation coefficient can be calculated as

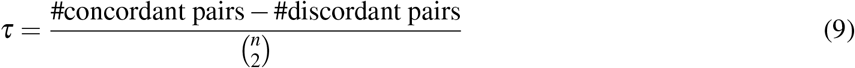

## Notes

### Competing Interest Statement

The authors have declared no competing interest.

## References

1. Haghverdi, L., Büttner, M., Wolf, F. A., Buettner, F. & Theis, F. J. Diffusion pseudo-time robustly reconstructs lineage branching. Nat. Methods 13, 845–848 (2016).

2. Street, K. et al. Slingshot: cell lineage and pseudo-time inference for single-cell transcriptomics. BMC Genomics 19, 477 (2018).

3. Tritschler, S. et al. Concepts and limitations for learning developmental trajectories from single cell genomics. Development 146 (2019).

4. La Manno, G. et al. RNA velocity of single cells. Nature 560, 494–498 (2018).

5. Bergen, V., Lange, M., Peidli, S., Wolf, F. A. & Theis, F. J. Generalizing RNA velocity to transient cell states through dynamical modeling. Nat. Biotechnol. (2020).

6. Saelens, W., Cannoodt, R., Todorov, H. & Saeys, Y. A comparison of single-cell trajectory inference methods. Nat. Biotechnol. (2019).

7. Zhang, X., Xu, C. & Yosef, N. Simulating multiple faceted variability in single cell RNA sequencing. Nat. Commun. 10, 2611 (2019).

8. Zappia, L., Phipson, B. & Oshlack, A. Splatter: simulation of single-cell RNA sequencing data. Genome Biol. 18, 174 (2017).

9. Vieth, B., Ziegenhain, C., Parekh, S., Enard, W. & Hellmann, I. powsimr: power analysis for bulk and single cell RNA-seq experiments. Bioinformatics 33, 3486–3488 (2017).

10. Cannoodt, R., Saelens, W., Deconinck, L. & Saeys, Y. dyngen: a multi-modal simulator for spearheading new single-cell omics analyses (2020).

11. Dibaeinia, P. & Sinha, S. SERGIO: A Single-Cell expression simulator guided by gene regulatory networks. Cell Syst (2020).

12. Peccoud, J. & Ycart, B. Markovian modeling of Gene-Product synthesis. Theor. Popul. Biol. 48, 222–234 (1995).

13. Munsky, B., Neuert, G. & van Oudenaarden, A. Using gene expression noise to understand gene regulation. Science 336, 183–187 (2012).

14. Zeisel, A. et al. Coupled pre-mRNA and mRNA dynamics unveil operational strategies underlying transcriptional responses to stimuli. Mol. Syst. Biol. 7, 529 (2011).

15. Felsenstein, J. Inferring phylogenies (Sinauer Associates, 2004).

16. Zhang, Z. & Zhang, X. Inference of multiple trajectories in single cell RNA-seq data from RNA velocity (2020).

